# Barbadin selectively modulates FPR2-mediated neutrophil functions independent of receptor endocytosis

**DOI:** 10.1101/2020.04.30.070011

**Authors:** Martina Sundqvist, André Holdfeldt, Shane C. Wright, Thor C. Møller, Esther Siaw, Karin Jennbacken, Henrik Franzyk, Michel Bouvier, Claes Dahlgren, Huamei Forsman

## Abstract

Formyl peptide receptor 2 (FPR2), a member of the family of G protein-coupled receptors (GPCRs), mediates neutrophil migration, a response that has been linked to β-arrestin recruitment. β-Arrestin regulates GPCR endocytosis and can also elicit non-canonical receptor signaling. To determine the poorly understood role of β-arrestin in FPR2 endocytosis and in NADPH-oxidase activation in neutrophils, Barbadin was used as a research tool in this study. Barbadin has been shown to bind the clathrin adaptor protein (AP2) and thereby prevent β- arrestin/AP2 interaction and β-arrestin-mediated GPCR endocytosis. In agreement with this, AP2/β-arrestin interaction induced by an FPR2-specific agonist was inhibited by Barbadin. Unexpectedly, however, Barbadin did not inhibit FPR2 endocytosis, indicating that a mechanism independent of β-arrestin/AP2 interaction may sustain FPR2 endocytosis. This was confirmed by the fact, that FPR2 also underwent agonist-promoted endocytosis in β-arrestin deficient cells, albeit at a diminished level as compared to wild type cells. Dissection of the Barbadin effects on FPR2-mediated neutrophil functions including NADPH-oxidase activation mediated release of reactive oxygen species (ROS) and chemotaxis reveled that Barbadin had no effect on chemotactic migration whereas the release of ROS was potentiated/primed. The effect of Barbadin on ROS production was reversible, independent of β-arrestin recruitment, and similar to that induced by latrunculin A. Taken together, our data demonstrate that endocytic uptake of FPR2 occurs independently of β-arrestin, while Barbadin selectively augments FPR2-mediated neutrophil ROS production independently of receptor endocytosis. Given that Barbadin binds to AP2 and prevents the AP2/β-arrestin interaction, our results indicate a role for AP2 in FPR2-mediated ROS release from human neutrophils.

## Introduction

Neutrophils, the most abundant leukocytes in human peripheral blood, form the frontline of our innate host immune defense, and are rapidly recruited from the circulation to damaged or infected body tissues, where they contribute to bacterial clearance and tissue repair [1-4]. The formyl peptide receptor 2 (FPR2), belonging to the family of G protein-coupled receptors (GPCRs), regulates directional neutrophil migration (chemotaxis), granule secretion (degranulation), formation of F-actin filaments (through polymerization of G-actin), and activation of the reactive oxygen species (ROS) producing NADPH-oxidase [3, 5, 6]. FPR2 recognizes not only N-formyl peptides of both bacterial and host mitochondrial origin, but also a large number of ligands belonging to different chemical classes [5-7]. Depending on the agonists examined, the signals generated by FPR2 mediate primarily pro-inflammatory activities, yet anti-inflammatory activities have also been described [8-11]. Based on the important roles of FPR2 in host defense and regulation of inflammation, this GPCR has been considered as a promising drug target for inflammatory conditions including cardiovascular diseases [12, 13]. FPR2 transduces signaling primarily through heterotrimeric G_i/o_ proteins and the dissociated Gβγ subunits trigger activation of phospholipase C (PLC) associated with an increase in intracellular Ca^2+^ [5, 7]. For most GPCRs, recruitment of β-arrestin desensitizes the agonist-occupied receptor at the plasma membrane, thereby terminating G protein-dependent signaling. In addition, recruitment of β-arrestin initiates receptor endocytosis/internalization and activation of non-canonical signaling [14]. The precise role of β-arrestin in regulating FPR2 signaling and function is not known, but it is clear that some FPR2 agonists, but not all, are capable of promoting recruitment of β-arrestin [5]. We have previously proposed that the dynamic transfer of agonist-occupied FPR2 from an active signaling state to a non-signaling state (i.e., receptor desensitization) is not directly associated with the ability of the receptor-agonist complex to recruit β-arrestin [9], but appears to involve receptor coupling to the actin cytoskeleton [11, 15-18]. Nevertheless, the ability to recruit β-arrestin is of functional importance also in neutrophils as demonstrated by the finding that FPR2 agonists that are biased away from β-arrestin recruitment induce ROS production without concomitant induction of chemotactic migration [9-11].

In the present study, we investigated the role of β-arrestin in regulating FPR2 endocytosis and the NADPH-oxidase activation in neutrophils by using Barbadin as a tool compound. Barbadin has been shown to bind the clathrin adaptor protein AP2 and by that prevent β-arrestin/AP2- mediated endocytosis of agonist-occupied GPCRs [19, 20]. Barbadin does not affect β-arrestin recruitment per se or internalization of GPCRs that rely on a β-arrestin/AP2-independent mechanism [19]. The data obtained show that (i) Barbadin did not inhibit endocytosis of agonist-occupied FPR2 despite its inhibitory effect on AP2/β-arrestin interaction, (ii) agonist-promoted FPR2 endocytosis was somewhat diminished in β-arrestin deficient cells and (iii) Barbadin both potentiates and resensitizes FPR2-mediated ROS production in neutrophils. Moreover, the potentiating effect by Barbadin on FPR2-mediated ROS production occurred regardless of if the agonist used had displayed a strong or very poor ability to recruit β-arrestin. In summary, we propose that FPR2 endocytosis can occur through a β-arrestin/AP2- independent mechanism and that Barbadin affects FPR2-mediated NADPH-oxidase activation independently of β-arrestin recruitment and receptor endocytosis.

## Material and Methods

### Ethics Statement

This study includes blood taken from healthy donors or from buffy coats obtained from the blood bank at Sahlgrenska University Hospital, Gothenburg, Sweden. According to the Swedish legislation section code 4§ 3p SFS 2003:460, no ethical approval was needed since the blood/buffy coats were provided anonymously.

### Chemicals

Barbadin was from MolPort (Riga, Latvia). Dextran and Ficoll-Paque were from GE-Healthcare Bio-Science (Piscataway, NJ, USA). Horseradish peroxidase (HRP), superoxide dismutase (SOD), phorbol 12-myristate 13-acetate (PMA) and Pefabloc protease inhibitor cocktail was from Roche Diagnostic (Mannheim, Germany). The hexapeptide WKYMVM and the F2Pal_10_ pepducin were acquired from CASLO Laboratory (Lyngby, Denmark). Isoluminol, luminol, latrunculin A, bovine serum albumin (BSA), poly-D-lysine, dimethyl sulfoxide (DMSO), Sigmacote, fluorescein-O’-acetic acid, and ATP were from Sigma (St. Louis, MO, USA), while catalase was from Worthington Biochemical Corporation (Lakewood, NJ, USA). Paraformaldehyde (PFA), Fura-2-acetoxymethyl ester (Fura-2AM), Dulbecco’s modified Eagle medium (DMEM), GlutaMAX, Hank’s balanced salt solution (HBSS), Lipofectamine 2000, Pluronic F-68 and Dulbecco’s phosphate-buffered saline (DPBS), Alexa Fluor (AF) 488- conjugated goat-anti-mouse antibody and AF647-conjugated phalloidin were from Thermo Fisher Scientific (Waltham, MA, USA). The rabbit anti-human AP2 complex subunit alpha-1 antibody was from Abcam (Cambridge, UK) while the goat anti-rabbit HRP-conjugated antibody was from Agilent Dako (Santa Clara, CA, USA). BD Cytofix/Cytoperm solution, the APC conjugated anti-human-CD11b antibody and yeast particles were from Becton Dickinson (San Jose, CA, USA), while the unconjugated mouse-anti-human-FPR2 antibody was from Santa Cruz (Dallas, TX, USA). Triton X-100 was from Merck (Darmstedt, Germany), the clarity western ECL substrate from Bio-Rad Laboratories (Hercules, CA, USA), and Tag-lite SNAP-Lumi4-Tb from Cisbio Bioassays (Codolet, France). Coelenterazine h and coelenterazine 400a were from Nanolight Technologies (Pinetop, AZ, USA). Compound 14 (i.e., Lau-[(S)-Aoc]-Lys-βNrpe-[Lys-βSNPhe]_5_-NH_2_, here denoted Cmp 14) and the FPR2- selective peptidomimetic antagonist (RhB-(Lys-βNPhe)_6_-NH_2_) were synthesized as described previously (structures are shown in Fig 1, [10, 21, 22]).

**Figure 1.**
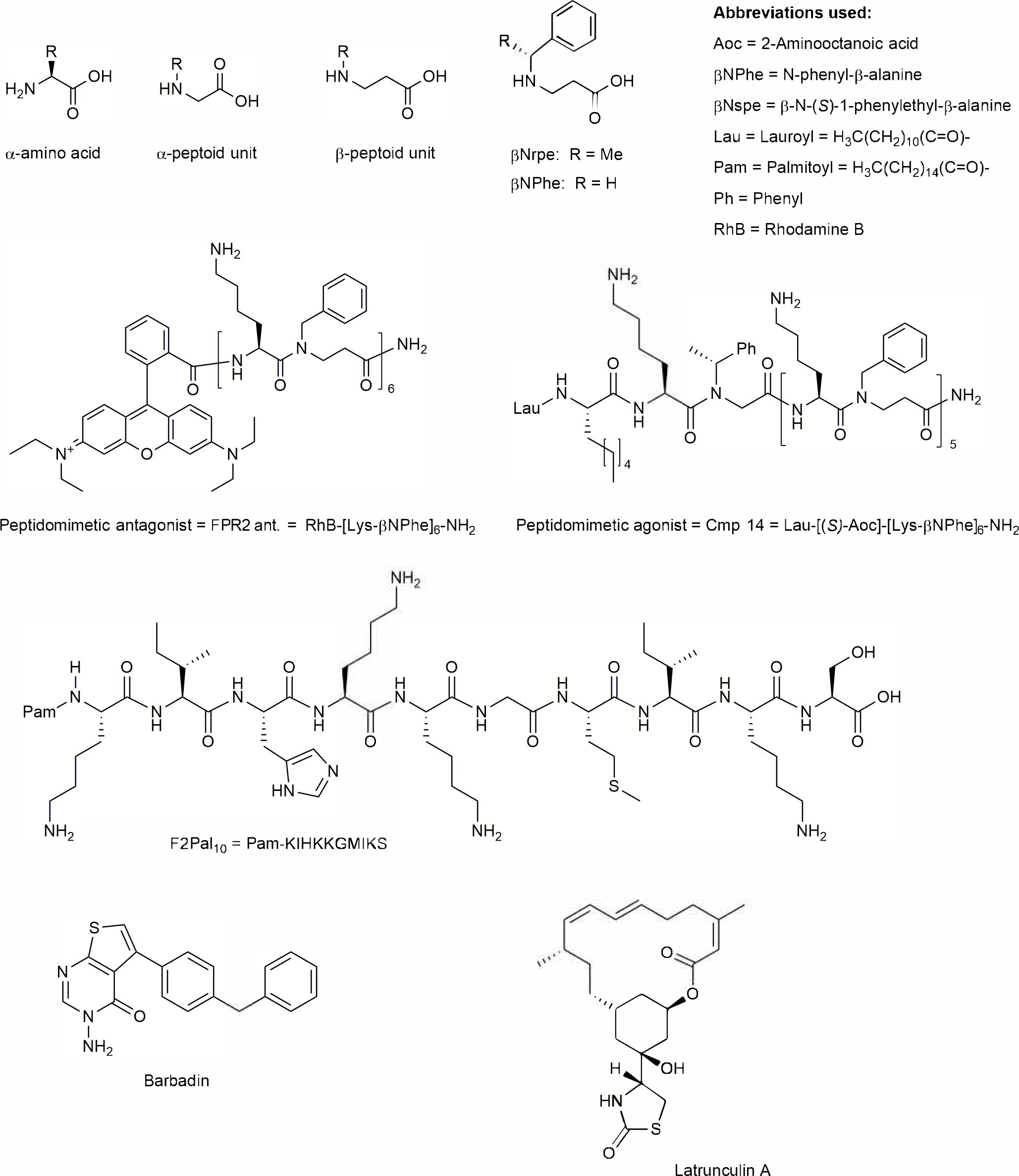
Chemical structures of compounds investigated in the study.

All reagents were dissolved in DMSO, before further dilution in Krebs-Ringer phosphate buffer (KRG, pH 7.3; 120 mM NaCl, 5 mM KCl, 1.7 mM KH_2_PO_4_, 8.3 mM NaH_2_PO_4_ and 10 mM glucose) supplemented with Ca^2+^ (1 mM) and Mg^2+^ (1.5 mM). Compounds used in TR-FRET experiments were further diluted in HBSS supplemented with 1 mM CaCl_2_, 1 mM MgCl_2_, 0.01% Pluronic F-68, and 20 mM HEPES, pH 7.4 (see time-resolved förster resonance energy transfer (TR-FRET) assay). For BRET experiments, dilution series with WKYMVM and F2Pal_10_ were added to HEK 293 cells to arrive at a final concentration of 1% DMSO whereas Compound 14 was diluted in Tyrode’s buffer (see Bioluminescence resonance energy transfer (BRET) assays).

### Plasmid DNA constructs

Plasmids encoding FPR2, V2R-*R*lucII, β-arrestin-2-*R*lucII, AP2-YFP, rGFP-CAAX, rGFP-FYVE have been previously described [13, 19, 23]. FPR2-*R*lucII was generated as follows: SP- FPR2 was PCR-amplified from pcDNA 3.1(+) SP-hFPR2 with T7 forward (5’- taatacgactcactataggg-3’) and SP-FPR2-*R*lucII Gibson reverse (5’- CTGCTCGGGGTCGTACACCTTGCTTGTCATGGTGGCTCTAGCCGGTGGATCCCCCGGTTTCATAGCCTGCAGCTCTGTTTCGGCAGG-3’) primers. *R*lucII was PCR-amplified from pcDNA3.1(+) PKC-BRET^1^ sensor with *R*lucII forward (5’-atgacaagcaaggtgtacgaccccgag- 3’) and *R*lucII EcoRI Gibson reverse (5’- CGCCACTGTGCTGGATATCTGCAGAATTCACTGTTCGTTCTTAAGCACTCTTTCCA C-3’) primers. The PCR fragments, SP-FPR and *R*lucII, were subcloned by Gibson assembly in pcDNA3.1 Zeo(+) digested with NheI+EcoRI. All plasmid constructs were verified by Sanger sequencing.

### Isolation of human peripheral blood neutrophils

Neutrophils from healthy donors were isolated from peripheral blood or buffy coats by dextran sedimentation, Ficoll-Paque gradient centrifugation and hypotonic lysis of remaining erythrocytes [24, 25]. The cells were suspended in KRG after isolation and stored on ice until use.

### Immunoblotting of AP2

Isolated neutrophils (1 × 10^7^/mL) were mixed 1:1 with reducing sample buffer supplemented with pefabloc (100 mM) and Triton X-100 (0.1%). Samples were then heated for 5 min at 90°C, centrifuged (16100 × *g*) for 15 s and resolved by SDS-PAGE. Proteins were transferred to nitrocellulose membranes, washed for 5 min (all washing steps were performed with PBS/0.05% Tween20) before blocked at room temperature (RT) for 1 h in PBS containing skimmed milk (5%)/Tween20 (0.05%). Membranes were then washed (3 x 5 min), prior to incubation with antibodies (all diluted in PBS/0.05% Tween20/5% BSA). Membranes were first incubated at RT for 2 h with a rabbit anti-human AP2 complex antibody (diluted 1:1000), followed by washing (3 × 5 min), prior to incubation with an HRP-conjugated goat anti-rabbit antibody (diluted 1:2000). After additional washing (3 x 5 min), ECL substrate was added according to manufacturer’s instructions after which chemiluminescence was analyzed in a Molecular Imager ChemiDoc XRS by Quantity One Software (Bio-Rad Laboratories, Hercules, CA, USA).

### Surface expression of FPR2 and CD11b (complement receptor 3; CR3) on human neutrophils

Neutrophil surface expression of FPR2 or CD11b was examined by flow cytometry. Neutrophils (5 × 10^6^/mL) were equilibrated for 20 min at 37°C, and then stimulated with either KRG (control), an FPR2 agonist or with barbadin with or without an FPR2 agonist in the presence of catalase (2000 Units/mL) to avoid potential agonist inactivation via oxidation. Cells (100 µL) were withdrawn at different time points after stimulation and fixed (20 min, 4°C) with PFA (2%), before washing and staining (30 min, 4°C) with an unconjugated mouse-anti-human FPR2 antibody (1:40 dilution), followed by washing and staining (30 min, 4°C) with an AF488- conjugated goat-anti-mouse antibody (1:400 final dilution). For CD11b surface expression, neutrophils (5 × 10^6^/mL) were incubated at 37°C in the absence or presence of Barbadin (10 µM; 20 min) or the positive control fMLF (10 nM; 10 min) prior to staining with an APC conjugated CD11b antibody (1:20 final dilution; 30 min at 4°C).

After staining, the cells were washed with PBS (10 min, 300 x *g*), resuspended and a minimum of 10,000 neutrophils (gated based on the forward scatter; size versus side scatter; density) per sample were collected on an Accuri C6 flow cytometer (BD Biosciences, Sparks, MD, USA). The surface expression of FPR2 and CD11b was determined by the geometric mean fluorescence intensity (MFI) as analyzed by FlowJo Software Version 10.3 (Tree star Inc., Ashland, OR, USA).

### Time-resolved Förster resonance energy transfer internalization assay

The plasmid encoding human FPR2 with an N-terminal SNAP-tag was obtained from Genscript (Piscataway, NJ, USA); it was constructed by replacing GLP1R in the previously described pcDNA3.1(+)-Flag-SNAP-GLP1R plasmid [26] with human FPR2. HEK 293A cells (Thermo Fisher Scientific) were cultured in DMEM with 4.5 g/L D-glucose and GlutaMAX supplemented with 10% (vol/vol), dialyzed fetal bovine serum and 100 units/mL Penicillin-Streptomycin at 37°C in a humidified 5% CO_2_ atmosphere. Prior to plating cells, white Falcon 96-well plates (Corning Inc., Corning, NY, USA) were siliconized with Sigmacote for 10-15 min, washed two times with DPBS and dried before coating with poly-D-lysine overnight. Cells (5 × 10^4^ cells/well) were transfected in the coated assay plate with 3 ng/well SNAP-FPR2, 37 ng/well pcDNA3.1(+) and 0.1 µL/well Lipofectamine 2000 by using a reverse transfection protocol. Real-time internalization was measured 24 h after transfection by using a previously described method [26]. SNAP-tagged receptors at the cell surface were labeled with 100 nM Tag-lite SNAP-Lumi4-Tb (donor) for 60 min at 37°C in assay buffer (HBSS buffer supplemented with 1 mM CaCl_2_, 1 mM MgCl_2_, 0.01% Pluronic F-68, and 20 mM HEPES, pH 7.4). Subsequently, cells were washed two times with assay buffer, and then 74 µL of fluorescein-O’-acetic acid (acceptor) diluted in assay buffer and 0.75 µL of Barbadin diluted in DMSO were added. After incubation (30 min, 37°C), 25 µL of WKYMVM or F2Pal_10_ (Pam- KIHKKGMIKS; structure is shown in Fig 1) diluted in assay buffer was added. Final concentrations were 50 µM fluorescein-O’-acetic acid, 10 µM Barbadin, 1 µM WKYMVM and 10 µM F2Pal_10_. Donor and acceptor emission were measured at 3 min intervals (during a total of 66 min) at 37°C with an EnVision 2014 Multilabel Reader (PerkinElmer, Waltham, MA, USA) using a 340/60 nm excitation filter and 520/8 nm (acceptor) and 615/8.5 nm (donor) emission filters. Donor/acceptor ratios were plotted to assess receptor internalization. All buffers and solutions were preheated to 37°C. Tubes and tips for dilution and addition of Barbadin were coated with Sigmacote.

### Bioluminescence resonance energy transfer assays

HEK 293 cells (ATCC, Manassas, VA, USA) were propagated in plastic flasks and on 6-well plates according to the manufacturer’s instructions. HEK 293 cells with targeted deletion of ARRB1 and ARRB2 (β-arrestin knockouts; ΔβARR) were derived, authenticated and propagated as previously described [27]. Both cell lines were cultured in DMEM supplemented with 10% FBS and 1% penicillin/streptomycin. For BRET experiments using Barbadin, cells were transfected in suspension with up to 1.0 μg of plasmid DNA using linear polyethyleneimine (PEI; MW 25,000) to a cell density of 350,000 cells/mL and grown on 6- well plates. After a 48 h incubation, cells were washed once with Tyrode’s buffer [140 mM NaCl, 2.7 mM KCl, 1 mM CaCl_2_, 12 mM NaHCO_3_, 5.6 mM D-glucose, 0.5 mM MgCl_2_, 0.37 mM NaH_2_PO_4_, 25 mM HEPES (pH 7.4)] supplemented with 0.1% BSA, harvested by trituration, and transferred to opaque white 96-well plates pre-coated with Sigmacote. The cells were incubated in the presence of Barbadin (100 μM) for 30 min prior to stimulation with agonist. BRET^1^ was subsequently measured after the addition of coelenterazine h (5 μM). For all other BRET experiments, cells were transfected in suspension with up to 1.0 μg of plasmid DNA and plated to a cell density of 350,000 cells/ml in white 96-well plates. After agonist stimulation, BRET^2^ was measured following the addition of coelenterazine 400a (5 μM). For receptor internalization experiments, cells were transfected with GPCR-*R*lucII and rGFP-CAAX or rGFP-FYVE in a (1:120) and (1:300) ratio for V_2_R and FPR2 respectively. For β- arrestin-2/AP2 interaction experiments, cells were transfected with FPR2, β-arrestin-2-*R*lucII and AP2-YFP in a (40:1:120) ratio. For β-arrestin-2 recruitment experiments, cells were transfected with FPR2, β-arrestin-2-*R*lucII and rGFP-CAAX in a (10:1:60) ratio. Cells were stimulated for 15 min with agonist before BRET measurements with the exception of the β- arrestin-2 recruitment where cells were stimulated for 5 min.

### Neutrophil NADPH-oxidase activity

NADPH-oxidase activity was determined by using isoluminol/luminol-enhanced chemiluminescence (CL) [5] in a six-channel Biolumat LB 9505 (Berthold Co., Wildbad, Germany). Polypropylene tubes containing a 900 µL reaction mixture of 10^5^ neutrophils in KRG, isoluminol (2 × 10^−5^ M) and HRP (4 Units/mL), were equilibrated at 37°C for 5 min before addition of 100 µL of stimulus. For latrunculin A or Barbadin dilution experiments, neutrophils (10^7^/mL) were pre-incubated at 37°C with latrunculin A (25 ng/mL) or Barbadin (10 µM) in a separate tube and then 10 µL cell suspension were transferred to polypropylene tubes containing 890 µL 37°C pre-heated reaction mixture (to obtain 10^5^ cells in the assay system) of KRG, HRP and isoluminol in the absence or presence of latrunculin A (25 ng/mL) or Barbadin (10 µM) followed by agonist stimulation (100 µL). For phagocytosis-induced intracellular NADPH-oxidase activity, yeast particles (1 × 10^8^/mL) were opsonized in 25% normal human serum (30 min, 37°C) followed by washing and dilution in KRG to a concentration of 5 × 10^7^ yeast particles/mL. Opsonized yeast solution (100 µL) was added to polypropylene tubes containing 900 µL reaction mixture containing 5 × 10^5^ neutrophils in KRG, luminol (2 × 10^−5^ M), superoxide dismutase (50 Units/mL) and catalase (2000 Units/mL) in the absence and presence of Barbadin (10 µM) or latrunculin A (25 ng/mL) that had been equilibrated at 37°C for 5 min; multiplicity of infection (MOI): 10:1. The light emission of yeast phagocytosis-induced ROS production is expressed as Mega counts per min (Mcpm).

### F-actin polymerization

Neutrophils (10^7^/mL) were equilibrated for 5 min in the absence and presence of Barbadin (10 µM) at 37°C followed by agonist stimulation. After 10 s of treatment, 100 μL cell suspension were transferred to ice-cold fixation/permeabilization solution (BD Cytofix/Cytoperm solution, 0.5 mL) and incubated on ice for 20 min. The cells were then washed twice with BD wash buffer before staining with AF647- conjugated phalloidin (30 min, 4°C). A minimum of 10,000 gated neutrophils (forward scatter; size versus side scatter; density) per sample were collected on an Accuri C6 flow cytometer (BD Biosciences, Sparks, MD, USA). The AF647- conjugated phalloidin intensity was determined by the geometric mean fluorescence intensity (MFI) as analyzed by FlowJo Software Version 10.3 (Tree star Inc., Ashland, OR, USA).

### Measurements of the transient rise in intracellular calcium [Ca^**2+**^]_**i**_

Neutrophils were loaded with Fura-2-AM (5 µM) (30 min, RT, in darkness) before washing and resuspension in KRG. Measurements of the transient rise in [Ca^2+^]_i_ were carried out at 37°C by using a PerkinElmer LC50 fluorescence spectrophotometer with excitation wavelengths of 340 nm and 380 nm, an emission wavelength of 509 nm, and slit widths of 5 nm and 10 nm. The transient rise in intracellular calcium is presented as the fluorescence intensities for both the excitation wavelengths (340 and 380 nm) as measured in parallel. For reactivation experiments (Fig 6C), isoluminol (2 × 10^−5^ M) and HRP (4 Units/mL) were included in the assay system to avoid agonist inactivation by the radicals generated through the myeloperoxidase (MPO)-H_2_O_2_ enzyme system [28].

**Figure 2.**
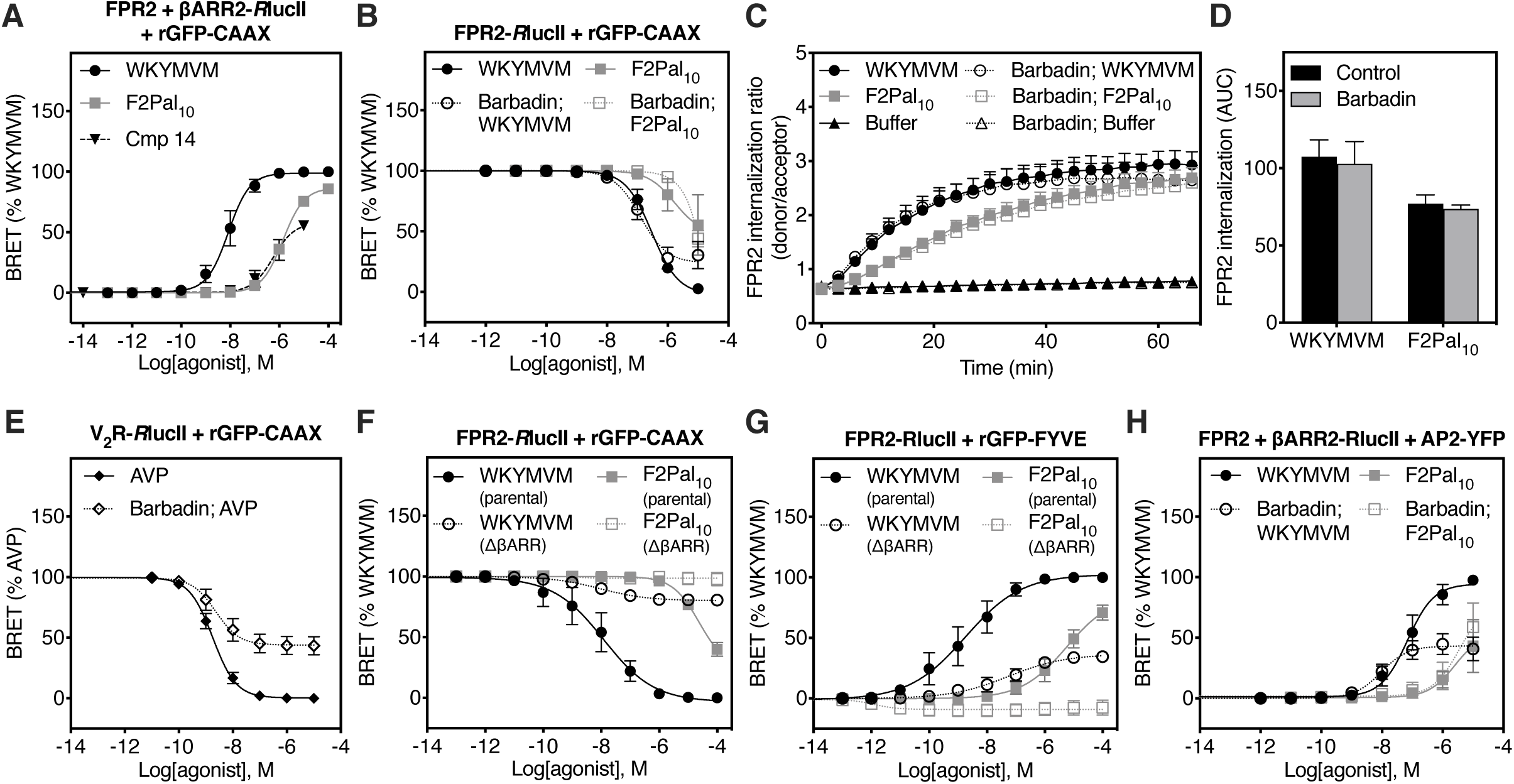
Barbadin does not affect FPR2 internalization in FPR2-overexpressing HEK 293 cells. (**A**) ebBRET experiments monitoring β-arrestin recruitment (mean ± SEM, n = 3-4 independent experiments). (**B, E**) ebBRET experiments monitoring receptor internalization. HEK 293 cells expressing either FPR2 (**B;** mean ± SEM, n = 2-6 independent experiments) or V_2_R (**E;** mean ± SEM, n = 4 independent experiments) were incubated with or without Barbadin (100 μM) for 30 min and stimulated for 15 min with agonist. (**C-D**) Time-resolved Förster resonance energy transfer internalization assay monitoring FPR2 internalization. FPR2-overexpressing HEK 293 cells were pre-incubated without (control) or with Barbadin (10 μM) for 5 min prior to stimulation with WKYMVM (1 μM), F2Pal_10_ (10 μM) or buffer. The amount of FPR2 internalization, represented as the area under the curve (AUC), was compared for each FPR2 agonist in the absence or presence of Barbadin (mean + SEM, n = 3-4 independent experiments). **(F-G)** ebBRET experiments were carried out in parental HEK 293 or ΔβARR cells to monitor trafficking of FPR2 from the plasma membrane (**F;** mean ± SEM, n = 5-6 independent experiments) to early endosomes (**G;** mean ± SEM, n = 6-7 independent experiments) after agonist stimulation. (**H**) BRET between β-arrestin2 and AP2 after agonist stimulation of FPR2 in the presence or absence of Barbadin (100 μM) (mean ± SEM, n = 3-5 independent experiments).

**Figure 3.**
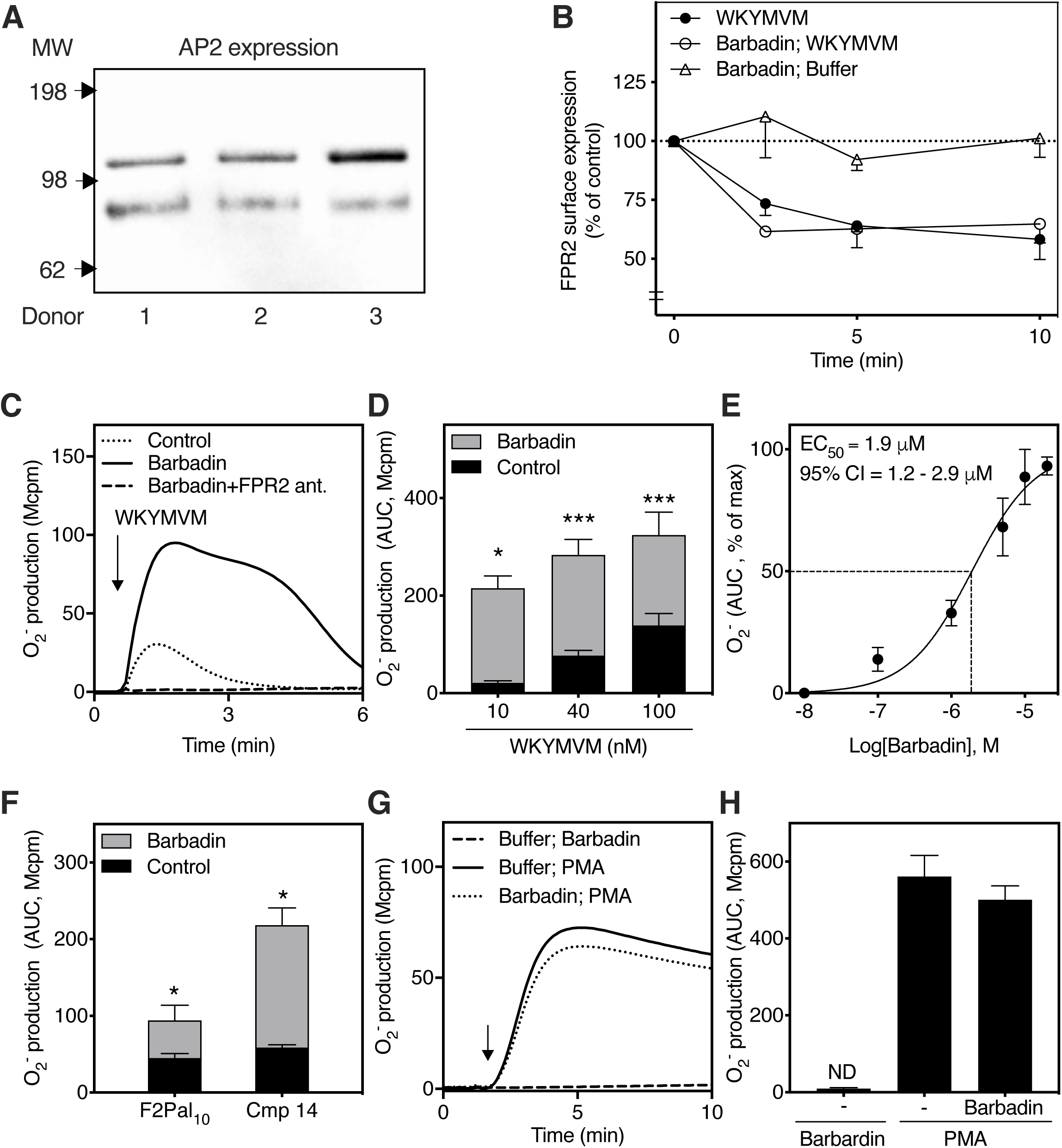
Barbadin augments FPR2-mediated ROS production in neutrophils. (**A**) Immunoblot of AP2 in naive neutrophils from three individual donors (each band represents one donor). (**B**) Neutrophils pre-incubated with and without Barbadin (10 μM) for 5 min at 37°C prior stimulation with buffer or WKYMVM (100 nM). The neutrophils were then stained with an anti-FPR2 antibody and surface-expressed FPR2 was and analyzed for geometric mean fluorescence intensity by flow cytometry. The data show the percentage of FPR2 expression as compared to cells prior to stimulation with WKYMVM (i.e., at time = 0 s; control; mean - SEM, n = 3 independent experiments). (**C**) Neutrophils were pre-incubated in the absence (control, dotted line) or presence of Barbadin (10 μM, solid line) or Barbadin (10 μM) together with the FPR2-selective peptidomimetic antagonist (RhB-[Lys-βNPhe]_6_-NH_2_, FPR2 ant.; 0.1 μM, dashed line) prior to WKYMVM stimulation (40 nM, indicated by an arrow). One representative trace out of three individual experiments is shown. (**D**) Neutrophils were pre-incubated in the absence (control, black bars) or presence of Barbadin (10 μM, grey bars) prior to stimulation with different concentrations of WKYMVM as indicated on the x- axis. The superimposed bar graph shows the amount of ROS induced (analyzed as the area under the curve; AUC) between neutrophils in the absence or presence of Barbadin for each WKYMVM concentration separately (mean + SEM; n = 3-4 independent experiments). (**E**) Barbadin dose-dependent priming effect for WKYMVM (40 nM). The EC_50_ value and 95% confidence interval (CI) was calculated based on the superoxide anion production as measured by the AUC (mean ± SEM, n = 3 independent experiments). (**F**) The superimposed bar graph shows the amount of ROS induced (analyzed as AUC) by stimulated neutrophils in the absence or presence of Barbadin, after addition of F2Pal_10_ (500 nM) or Cmp 14 (50 nM) separately (mean + SEM, n = 3 independent experiments). (**G**-**H**) Neutrophils were pre-incubated in the absence (control, solid line) or presence of Barbadin (10 μM, dotted line) prior to stimulation with PMA (50 nM, indicated by an arrow). Control cells stimulated with Barbadin (10 μM, dashed line) are also shown. The bar graph shows the AUC of superoxide anion production in the absence (control) or presence of Barbadin (mean + SEM, n = 3-8 independent experiments, ND; non detectable).

**Figure 4.**
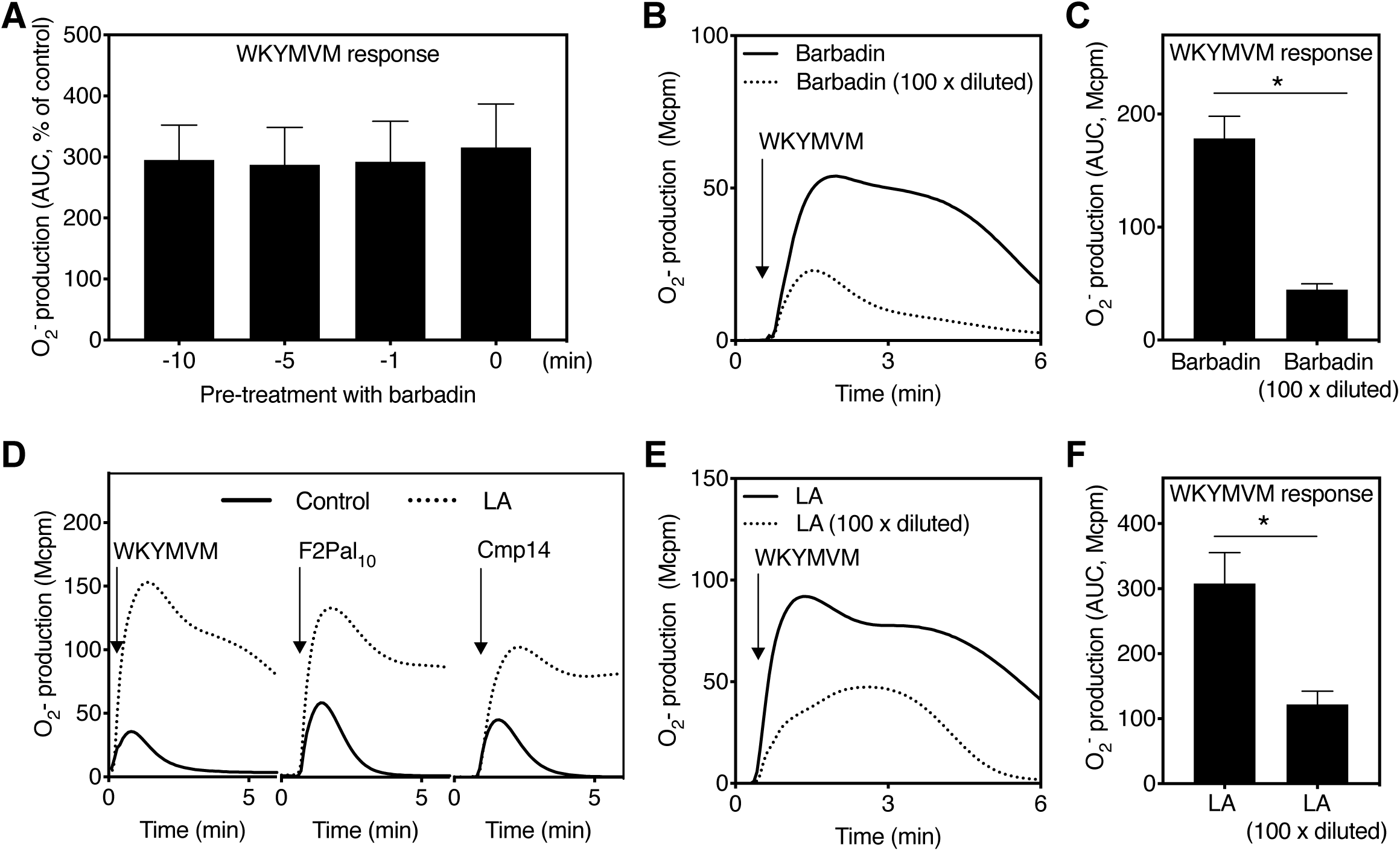
Barbadin-mediated priming of the NADPH-oxidase is reversible and resembles that of latrunculin A. (**A**) Neutrophils were pre-treated with Barbadin (10 μM) for different time points (indicated on the x-axis) prior to stimulation with WKYMVM (100 nM). The area under the curve (AUC) of the superoxide anion production was compared to the control response in the absence of Barbadin (mean + SEM, n = 3 independent experiments). (**B**-**C**) Neutrophils (10^7^/mL) were pre-incubated with Barbadin (10 μM) at 37°C for 5 min followed by a transfer of 10 μL cell suspension to chemiluminescence (CL) reaction tubes containing 990 μL CL reagents with Barbadin (10 μM) or buffer (Barbadin; 100-fold diluted). (**B**) The superoxide anion production was induced by WKYMVM (100 nM) and measured over time. (**C**) The bar graph shows the AUC of WKYMVM-induced superoxide anion production (mean + SEM, n = 3 independent experiments). (**D**) Neutrophils were pre-incubated in the absence (control) or presence of latrunculin A (LA; 25 ng/mL) for 5 min at 37°C prior to stimulation (indicated by an arrow) with WKYMVM (40 nM), F2Pal_10_ (500 nM) or Compound 14 (Cmp 14; 50 nM). One representative CL trace out of 3-5 individual experiments for each agonist is shown. (**E**-**F**) Neutrophils (10^7^/mL) were pre-incubated with LA (25 ng/mL) before 10 μL of the LA-treated cell suspension was transferred to tubes containing 990 μL CL reagents in the presence of LA (25 ng/mL) or buffer (LA; 100 x diluted). (**E**) Superoxide anion production was induced by WKYMVM (100 nM) and measured over time. (**F**) The bar graph shows the AUC of WKYMVM-induced superoxide anion production (mean + SEM, n = 3 independent experiments).

**Figure 5.**
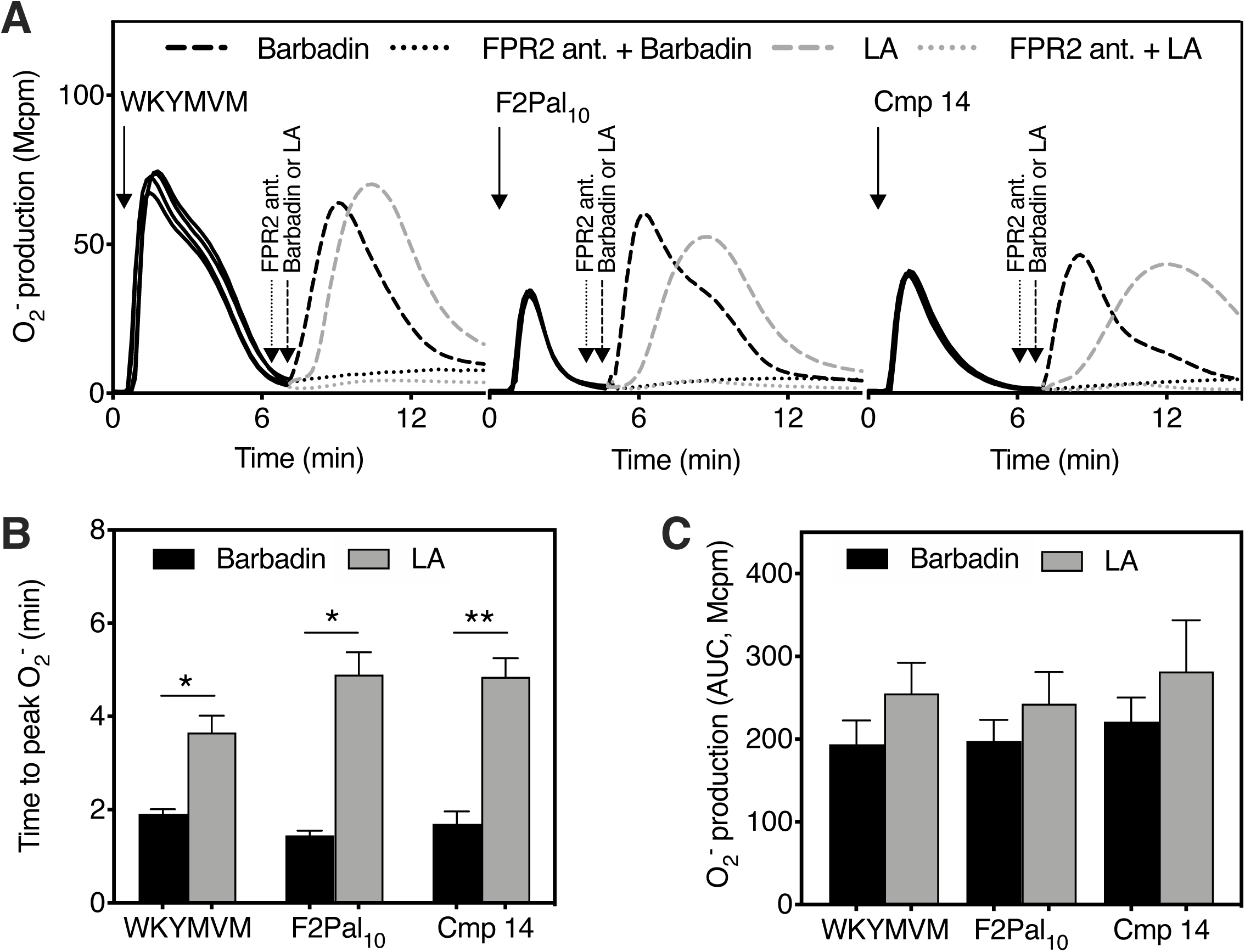
Barbadin triggers reactivation of desensitized FPR2 with a similar profile as latrunculin A. (**A**-**C**) Neutrophils were pre-incubated at 37°C for 5 min prior to stimulation with the FPR2 agonists (black solid lines; stimulation indicated by solid arrows): WKYMVM (100 nM), F2Pal_10_ (500 nM,) or Cmp 14 (50 nM) and measurement of superoxide anion release over time. Once the agonist induced response returned to basal levels, the neutrophils were treated with (dotted lines) and without (dashed lines) the FPR2-selective peptidomimetic antagonist (RhB- (Lys-βNPhe)_6_-NH_2_, FPR2 ant.; 0.1 μM) for 1 min (indicated by dotted arrows), prior to stimulation (indicated by dashed arrows) with Barbadin (10 μM; black dotted and dashed lines) or latrunculin A (LA, 25 ng/mL; grey dotted and dashed lines). (**A**) One out of three individual traces are shown for each agonist. (**B**) The time for the peak superoxide anion response induced after addition of Barbadin or LA was calculated and compared (mean + SEM, n = 3 independent experiments). (**C**) The area under the curve (AUC) of the superoxide anion production induced after addition of Barbadin or LA was calculated and compared (mean + SEM, n = 3 independent experiments).

**Figure 6.**
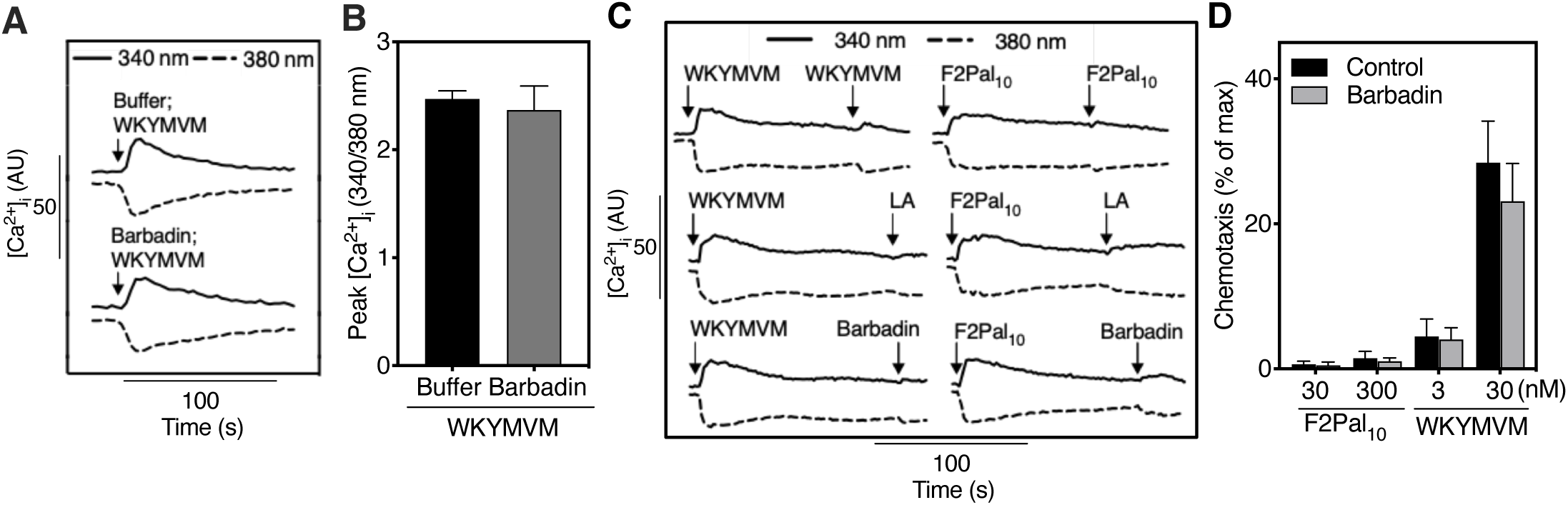
Barbadin lacks effect on FPR2-mediated transient rise in [Ca^**2+**^]_**i**_ **and chemotaxis**. (**A-B**) Neutrophils loaded with Fura-2AM were incubated in the absence (Buffer) or presence of Barbadin (10 μM), for 10 min at 37°C. Thereafter, the cells were stimulated with WKYMVM (5 nM) and the transient rise in cytosolic calcium ([Ca^2+^]_i_; y-axis), measured as the fluorescence intensities for both the excitation wavelengths (340 and 380 nm), was analyzed over time (seconds; s, x-axis). (**A**) One representative trace of the rise in [Ca^2+^]_i_ out of three individual experiments is shown. (**B**) The bar graph shows the peak [Ca^2+^]_i_, determined based on the ratio between 340 and 380 nm excitation wavelengths (mean + SEM, n = 3). (**C**) To mimic the protocol used for reactivation of desensitized neutrophils, Fura-2AM loaded cells were pre- warmed for 10 min at 37°C in the presence of isoluminol and HRP before stimulation with the FPR2 agonists WKYMVM (100 nM) or F2Pal_10_ (500 nM) and measurement of [Ca^2+^]_i_ over time. The cells showed only a low response to a second stimulation with (upper panel) the same agonist, (middle panel) latrunculin A (LA; 25 ng/mL) or (lower panel) Barbadin (10 μM). One representative trace out of three individual experiments are shown. (**D**) Neutrophil migration was measured by using a transwell migration system. For quantitative analysis, the migrated cells were lysed and the level of MPO was determined. All values were calculated by subtracting the MPO level obtained from the wells without any attractant. The bar graph (mean + SEM, n = 4-6 independent experiments) represents the fraction of cells that migrated towards F2Pal_10_ (30 and 300 nM) or WKYMVM (3 and 30 nM) in the absence (control) or presence of Barbadin (10 μM; added in both upper and lower chambers to prevent any gradient), shown as the percentage of cells recovered in the lower compartment as compared to the positive control (max; neutrophils added directly to the lower chambers).

### Chemotaxis

Neutrophils (2 × 10^6^ /mL) in KRG supplemented with BSA (0.3%) were loaded on top of a 3- µm pore size filter and allowed to migrate towards stimuli loaded into the bottom wells of a chemotaxis plate (ChemoTx, Neuro probe, UK) for 90 min at 37°C, 5% CO_2_. Barbadin was present in both upper and lower chambers to avoid any gradient effect. The migrated cells were quantified by measuring myeloperoxidase activity of cell lysates [11]. All migration (myeloperoxidase activity) values were subtracted from the level of migration without any attractant in the lower compartment (negative control), and the resulting data are presented as the number of cells recovered in the lower compartment, relative to the total number of cells applied to the migration system (neutrophils were added directly to the bottom well of the chemotaxis plate).

### Data analysis

Data analysis was performed using GraphPad Prism 8.1.0a (Graphpad Software, San Diego, CA, USA). Curve fitting was performed by non-linear regression using the sigmoidal dose-response equation (variable-slope). Independent BRET experiments were normalized to the span of reference compound (WKYMVM) as determined by non-linear regression analysis and pooled together. Data represent mean values ± SEM. Statistical analysis was performed using a paired Student’s *t*-test (Fig 2D; 3D, F and H; 4C and F; 5B and C; 6B and D and 7F) or a repeated measurement one-way ANOVA followed by Tukey’s multiple comparison test (Fig 4A, 7A and 7H). Statistically significant differences are indicated by \**p* < 0.05, \*\**p* < 0.01, \*\*\**p* < 0.001.

## Results

### Barbadin does not prevent agonist-induced internalization of FPR2 in receptor overexpressing HEK 293 cells

Barbadin (structure is shown in Fig 1) was recently described as a selective inhibitor of the machinery responsible for AP2/β-arrestin-dependent endocytic internalization of some GPCRs including the vasopressin receptor 2 (V_2_R), the β_2_-adrenergic receptor (β2AR) and the angiotensin II receptor type 1 (AT_1_R) [19]. To study the effect of Barbadin on agonist-induced FPR2 internalization, we used the peptide agonist WKYMVM and two lipopeptides, the peptidomimetic Cmp 14 and the pepducin F2Pal_10_ that are functionally biased (structures are shown in Fig 1). All three agonists (WKYMVM, Cmp 14 and F2Pal_10_) are potent in inducing an FPR2-mediated transient rise in [Ca^2+^]_i_, ERK1/2 phosphorylation, and ROS production in human neutrophils, but Cmp 14 and F2Pal_10_ are biased away from β-arrestin recruitment and chemotaxis [9-11]. By using an enhanced bystander bioluminescence resonance energy transfer (ebBRET) assay system, we here confirmed that Cmp 14 and F2Pal_10_ are poor inducers of β- arrestin recruitment (Fig 2A). Both WKYMVM and F2Pal_10_ triggered FPR2 internalization but Barbadin lacked effect on FPR2 internalization as determined by ebBRET (Fig 2B). The inability of Barbadin to inhibit agonist-triggered removal of FPR2 from the HEK 293 cell surface was also evident when time resolved-Förster Resonance Energy Transfer assay (TR-FRET) [26] was used to study FPR2 internalization (Fig 2C, D). These data clearly show that internalization of FPR2 induced by agonists is unaffected by the presence of Barbadin.

In contrast to FPR2, and in agreement with previously published data [19], Barbadin reduced V_2_R internalization as measured by ebBRET in HEK 293 cells that had been transiently transfected with donor-tagged V_2_R (V_2_R-*R*lucII) and an acceptor anchored to the membrane (rGFP-CAAX) (Fig 2E). Although Barbadin did not appear to block agonist-induced FPR2 internalization, ebBRET experiments investigating FPR2 trafficking from the plasma membrane (rGFP-CAAX) to early endosomes (rGFP-FYVE) revealed that the internalization was largely (but not completely for WKYMVM) dependent on the presence of β-arrestin1/2 as illustrated by the reduced responses in HEK 293 cells devoid of β-arrestin1/2 (ΔβARR, Fig 2F, G). In order to confirm that Barbadin did indeed block the interaction between β-arrestin-2 and AP2, we performed BRET experiments to determine the WKYMVM and F2Pal_10_ induced interaction between the two proteins. Barbadin clearly blocked the WKYMVM-induced interaction between β-arrestin-2 and AP2, whereas the low level of interaction induced by F2Pal_10_ alone makes it hard to determine the ability/inability of Barbadin to inhibit its effect (Fig 2H). Taken together, these findings demonstrate that Barbadin does not affect agonist-triggered internalization of surface-exposed FPR2s and that a mechanism independent of β- arrestin/AP2 interaction can sustain WKYMVM-induced receptor endocytosis.

### Barbadin potentiates the FPR2-mediated generation of ROS in neutrophils

We next determined whether Barbadin affects agonist-induced internalization of FPR2 when it is endogenously expressed in human neutrophils. Although AP2 was clearly present in neutrophils (Fig 3A), the reduction in the number of surface-exposed FPR2s was identical in WKYMVM-activated control neutrophils (not treated with Barbadin) and in WKYMVM- activated cells treated with Barbadin (Fig 3B). Hence, these data strongly imply that Barbadin has no effect on agonist-induced FPR2 internalization in primary human neutrophils in which the receptor is endogenously expressed.

Neutrophils possess an electron transporting enzyme system, the neutrophil NADPH-oxidase, which upon activation generates ROS [5]. Despite the insensitivity of the agonist-triggered FPR2 internalization to Barbadin, pre-incubation of neutrophils with Barbadin substantially increased the neutrophil release of ROS upon WKYMVM stimulation (Fig 3C, D). Expectedly, the FPR2-selective antagonist RhB-(Lys-βNPhe)_6_-NH_2_ (structure is shown in Fig 1, where it is denoted as FPR2 ant.) completely blocked the neutrophil response (Fig 3C). Also, the priming effect (i.e., potentiation of ROS production) of Barbadin was evident at a concentration of the peptide agonist WKYMVM (10 nM) too low to trigger ROS release in non-primed neutrophils (Fig 3D). The priming effect was concentration-dependent, reaching its full effect at 10 μM of Barbadin with an EC_50_ value of ∼2 μM (Fig 3E). Similarly, Barbadin also significantly primed the ROS production induced by F2Pal_10_ and Cmp 14 (Fig 3F). The fact that F2Pal_10_ and Cmp 14 cannot induce β-arrestin recruitment at these concentrations ([9-11], Fig 2A), strongly suggests that the priming effect of Barbadin on FPR2-mediated ROS production is independent of the ability of the activating agonist to recruit β-arrestin. In addition, the ROS release following activation with PMA, a compound that bypasses receptors and directly activate protein kinase C (PKC), was unaffected by Barbadin (Fig 3G, H), which suggests that Barbadin lacks a direct effect on the ROS-producing NADPH-oxidase machinery. It is also important to note that activation of the NADPH-oxidase could not be triggered by Barbadin alone (Fig 3G, H).

Collectively, these results suggest that Barbadin primes neutrophils in their response to FPR2 agonists, and the increased NADPH-oxidase activation is regulated independently of FPR2 internalization and FPR2-induced β-arrestin recruitment.

### Barbadin resensitizes FPR2-desensitized neutrophils – a feature that resembles the function of the F-actin-disrupting agent latrunculin A

Barbadin treatment significantly primed FPR2 agonist-induced ROS production in human neutrophils and further characterization revealed that a very short interaction time between neutrophils and Barbadin was needed for Barbadin to exert its priming effect. In fact, the increase in neutrophil ROS production was the same when Barbadin and the FPR2 agonist WKYMVM were added simultaneously, as when the cells were incubated with Barbadin prior to addition of the activating peptide WKYMVM (Fig 4A). In addition, we found that the priming effect of Barbadin on neutrophils for increased WKYMVM response was also rapidly reduced when the Barbadin concentration was lowered significantly. Neutrophils first incubated for 5 min with an effective priming concentration of Barbadin (10 µM) and then transferred to two different ROS measurement reagents, one containing a new addition of Barbadin (10 µM) and the other not (the Barbadin concentration was thus lowered 100 x to 0.1 µM), followed by stimulation with WKYMVM. A significantly lower degree of priming effect was observed in cells that were exposed to a reduced concentration (from 10 µM to 0.1 µM) than cells incubated constantly to an effective priming concentration (10 µM) of Barbadin (Fig 4 B and C). This suggests that the neutrophil priming induced by Barbadin is a process that is reversible, unlike many other priming processes that involve an irreversible process of neutrophil granule secretion [17, 29, 30]. The effect of Barbadin on FPR2-mediated ROS production resembled the effect induced by the F-actin-disrupting agent latrunculin A. Barbadin and latrunculin A both prolonged and increased the magnitude of FPR2-mediated neutrophil ROS production as compared to the corresponding ROS production in non-treated control cells (Fig 4D). In addition, very similar to Barbadin, the priming effect of latrunculin A was rapidly reduced as shown when latrunculin A was removed prior to activation with the FPR2 agonist WKYMVM (Fig 4E, F), using the same dilution protocol, to show that the priming effect of Barbadin is reversible, as described above.

FPR2-induced ROS production is a process rapidly initiated after addition of an activating agonist, and within a time period of minutes, the ROS production is terminated and the cells are homologously desensitized. The desensitized cells are non-responsive to a second stimulation with FPR2 agonists but fully responsive to an FPR1 selective agonist or PMA [11, 31]. The desensitized state could be transferred to an active signaling state to produce ROS again (i.e., FPR2 resensitization or reactivation) when the cytoskeleton was disrupted by latrunculin A (Fig 5A). The ROS production induced by latrunculin A in FPR2-desensitized cells was completely abolished by the FPR2-selective antagonist RhB-(Lys-βNPhe)_6_-NH_2_ (Fig 5A). To determine the ability of Barbadin to resensitize desensitized FPR2, we reversed the order by which the sensitizer (Barbadin) and FPR2 agonist (WKYMVM) were added to the neutrophils. Barbadin was added to WKYMVM-activated neutrophils at a time point when ROS production had returned to a background level; interestingly, these FPR2-desensitized cells could be resensitized to produce ROS also by Barbadin, similar to the effect of latrunculin A (Fig 5A). This response was also inhibited by an FPR2-selective antagonist (Fig 5A), clearly demonstrating an FPR2-mediated neutrophil resensitization. Very similar results were obtained in FPR2-desensitized cells when F2Pal_10_ or Cmp 14 (at concentrations that could not recruit β- arrestin [9-11], Fig 2A) replaced WKYMVM as the agonist used to desensitize FPR2 (Fig 5A). Although the resensitization effects of latrunculin A and Barbadin appeared very similar, it should be noted that the lag phase before any ROS were generated was shorter when Barbadin was used for resensitization, and the time to reach maximal (peak) ROS production was also shorter as compared to the response following addition of latrunculin A (Fig 5B). However, the amounts of ROS produced during resensitization (as measured by the area under the curve) were comparable for Barbadin and latrunculin A (Fig 5C).

In summary, we show that Barbadin, similarly to latrunculin A, not only potentiates the ROS production induced by the different FPR2 agonists, but also resensitizes FPR2 signaling when added to FPR2 desensitized neutrophils.

### Barbadin lacks effect on FPR2-mediated transient rise in [Ca^**2+**^]_**i**_ **and chemotaxis**

Barbadin treatment potentiated and resensitized FPR2-mediated signaling leading to ROS production. To further investigate the effect of Barbadin on FPR2 signaling and function in neutrophils, we measured the transient rise in cytosolic calcium [Ca^2+^]_i_ mediated through FPR2. In contrast to the potentiating effect on ROS production, the rise in [Ca^2+^]_i_ induced by WKYMVM was not affected by Barbadin (Fig 6A, B). Similarly, resensitization by Barbadin of agonist-desensitized FPR2 leading to an activation of the ROS producing NADPH-oxidase, was not associated with a corresponding rise in [Ca^2+^]_i_ (Fig 6C).

At the functional level, we also observed a biased activity of Barbadin in favor of neutrophil ROS production over directional cell migration/chemotaxis. Neutrophil chemotaxis was measured by using a transwell migration system; neutrophils were placed on top of the filter and allowed to migrate towards different concentrations of FPR2 agonists that were placed in the bottom well of the chambers. Barbadin (10 μM) was added to both compartments, so that it was present in both the upper chamber together with the cells and the lower chamber containing the agonist. In line with earlier data [9, 11], WKYMVM, but not F2Pal_10_, triggered a chemotactic migration of neutrophils. Further, neutrophil chemotaxis towards WKYMVM was unaffected by Barbadin, and the inability of F2Pal_10_ to trigger chemotaxis was retained in the presence of Barbadin (Fig 6D). In summary, these data demonstrate that the effect of Barbadin in neutrophils is in favor of FPR2-mediated ROS over the rise in [Ca^2+^]_i_ and chemotaxis.

### Barbadin lacks a direct effect on the dynamic re-organization of the actin cytoskeleton and on mobilization of granule-localized receptors

The functional similarities between Barbadin and latrunculin A, both being priming agents affecting the response induced by FPR2 agonists and agents that resensitize desensitized FPR2, promoted us to examine whether Barbadin could directly affect the integrity of the actin cytoskeleton and granule secretion. WKYMVM induced a rapid polymerization of G-actin monomers into polymerized filamentous actin (F-actin) in human neutrophils (Fig 7A). However, the presence of Barbadin did not affect the formation of F-actin induced by WKYMVM (Fig 7A), indicating that Barbadin does not affect the integrity of the actin cytoskeleton.

**Figure 7.**
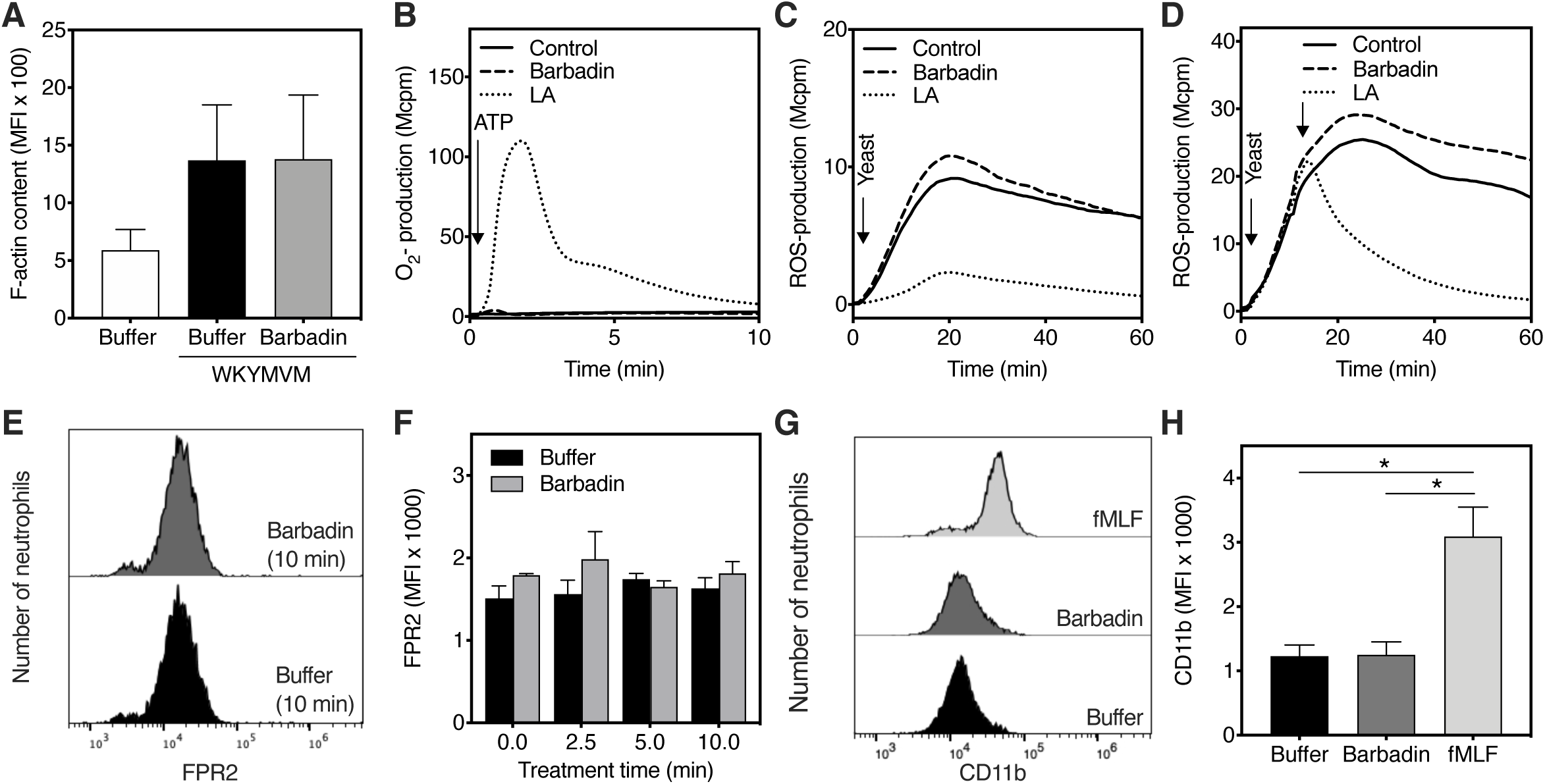
Barbadin does not interfere with actin cytoskeleton-dependent functions nor granule secretion process. (**A**) Neutrophils were pre-incubated at 37°C in the absence or presence of Barbadin (10 μM) for 5 min before being challenged with WKYMVM (10 nM), followed by further incubation at 37°C for 10 s. Thereafter, actin polymerization was stopped by simultaneous fixation/permeabilization followed by staining of the F-actin with AF647-conjugated phalloidin. The bar graph shows the geometric mean fluorescence intensity (MFI) obtained by flow cytometry analysis (mean + SEM; n = 4 independent experiments). (**B**) Neutrophils were pre-incubated at 37°C for 5 min in the absence (control) or presence of Barbadin (10 μM) or LA (25 ng/mL) prior to stimulation (indicated by an arrow) with ATP (50 μM), followed by measurement of superoxide anion production over time. One representative trace out of four individual experiments is shown. (**C-D**) Neutrophils were triggered with yeast (MOI 10) and measured for intracellular ROS-production over time. One representative trace out of three individual experiments is shown. (**C**) Neutrophils were pre-incubated at 37°C for 5 min in the absence (control) or presence of Barbadin (10 μM) or LA (25 ng/mL) prior to stimulation (indicated by an arrow). (**D**) Neutrophils were pre-incubated at 37°C for 5 min prior to stimulation. After ∼15 min stimulation, Barbadin (10 μM) or LA (25 ng/mL) was added (indicated by an arrow). **(E-H)** Analysis of FPR2 and CD11b surface expression was performed by flow cytometry. **(E-F)** Neutrophils were incubated in the absence (buffer) and presence of Barbadin (10 μM) at 37°C for different time points as indicated on the x-axis, prior fixation and staining with an anti-FPR2 antibody. (**G**-**H**) Neutrophils were incubated in the absence (buffer, 20 min) and presence of Barbadin (10 μM; 20 min) or fMLF (10 nM; 10 min) at 37°C prior staining with an anti-CD11b antibody. (**E, G**) Representative histograms are shown. (**F, H**) The bar graphs show the MFI of (**G**) FRP2 and (**H**) CD11b expression (mean + SEM; n = 3 independent experiments).

The observation that Barbadin does not directly interfere with the integrity of the actin cytoskeleton gained further support from the results obtained with Barbadin on two neutrophil responses previously found to be regulated by the actin cytoskeleton, i.e., the ATP receptor P2Y_2_R-mediated ROS production and the phagocytosis process [32, 33]. It is known that ATP upon binding to its neutrophil receptor P2Y_2_R, triggers ROS production provided that the F-actin structure is disrupted (Fig 7B). In line with this, latrunculin A treated neutrophils produce ROS when activated with ATP, but no such effect was obtained with Barbadin (Fig 7B). Activation of the ROS-generating NADPH-oxidase system during uptake (phagocytosis) of microbes is a process regulated by the cytoskeleton, and accordingly, latrunculin A inhibited the activation process (Fig 7C). However, Barbadin exerted no inhibitory effect, neither when added before addition of the phagocytosis prey nor when added during the ongoing activation process (Fig 7C, D). Taken together, these data show that Barbadin has no direct effects on basic neutrophil functions regulated by the actin cytoskeleton, supporting the conclusion that Barbadin lacks a general effect on the assembly of the ROS-producing oxidase.

Our data reveal that Barbadin is able to prime neutrophils for enhanced FPR2-mediated ROS production. Neutrophil priming is a well-known process both *in vitro* and *in vivo* [17, 29, 30, 34, 35], and an increased exposure of intracellular granule-localized receptors to the plasma membrane as a result of granule secretion has been suggested to be one of the main mechanisms that augments neutrophil NADPH-oxidase activity [17, 29, 30]. We next determined whether the receptor mobilizing effect with increased surface FPR2 expression could account for the Barbadin-induced priming effect. However, no increased surface exposure of FPR2 was induced by incubation of neutrophils with Barbadin for up to ten minutes (Fig 7E, F). In addition to FPR2 expression, the ability of Barbadin to upregulate the surface expression of CD11b (complement receptor 3; CR3), a marker protein stored in easily mobilized neutrophil granule compartments that can be mobilized to the surface by many secretagogues or priming agents [36] was investigated. However, similar to the FPR2 expression, Barbadin also lacked effect to upregulate CD11b on the plasma membrane, whereas a profound increase of CD11b surface expression was induced by the classical secretagogue fMLF (Fig 7G, H).

In summary, our data show that even though Barbadin affects FPR2 signaling in a way that resembles actin cytoskeleton-disrupting agents, Barbadin lacks direct effects on the re- organization of the actin cytoskeleton in neutrophils and on receptor mobilization from intracellular granule stores.

## Discussion

In the present study, we assessed the role of β-arrestin in endocytosis of FPR2 and in receptor down-stream functional responses in human neutrophils using Barbadin, an AP2-binding inhibitor that blocks the interaction between β-arrestin and AP2 and prevents agonist triggered endocytosis of many GPCRs [19]. Our data show that the AP2 protein targeted by Barbadin indeed is expressed in neutrophils, yet, Barbadin did not block FPR2 endocytosis. These results imply that FPR2 can be internalized through a β-arrestin/AP2-independent process, an assumption in line with the observation that only residual endocytosis of FPR2 occurs in cells lacking β-arrestin. Interestingly, Barbadin treatment potentiated FPR2-mediated ROS production and resensitization of FPR2-desensitized human neutrophils in a manner similar as an inhibitor of actin polymerization (i.e., latrunculin A). However, Barbadin did not interfere with other processes in neutrophils involving the actin cytoskeleton machinery. In addition, the potentiating effect of Barbadin on FPR2-mediated ROS production was found to involve biased functional/signaling as neither FPR2-promoted intracellular calcium mobilization nor chemotaxis was affected when AP2-binding was inhibited by Barbadin.

Previously, Barbadin has been shown to affect several GPCR-mediated functions, including hormone secretion mediated by gonadotropin-releasing hormone (GnRH) receptors [37], and uptake/entry of influenza A viruses facilitated by short chain fatty acid receptor 2 (FFAR2) signaling [20]. As described, Barbadin prevents AP2/β-arrestin-mediated receptor endocytosis, which has been deemed to be the canonical molecular mechanism behind the functional effects of this AP2 inhibitor. However, the data presented in the current study suggest that alternative endocytosis- and β-arrestin-independent mechanisms can mediate the effects by this AP2- binding inhibitor with regards to FPR2 expressing human neutrophils. Recent data infer that β- arrestin appears to be involved in non-canonical and endosomal signaling, besides playing roles in receptor desensitization and endocytosis [14, 38]. However, the exact functional role of β- arrestin in FPR2 signaling needs to be further investigated. It has been suggested that polymerized actin rather than β-arrestin constitutes the basis for physical separation of G proteins from activated FPRs, resulting in termination of signaling and receptor desensitization [15, 16, 18]. The role of the actin cytoskeleton in the regulation of GPCR signaling in neutrophils was originally defined by measurements of ROS generated by the phagocyte NADPH-oxidase. Involvement of the actin cytoskeleton in the termination/desensitization of FPR signaling became evident from experiments in which actin cytoskeleton disrupting agents prolong FPR signaling and have the ability to resensitize the desensitized receptors [11, 17, 18]. We now show that Barbadin lacks effect on agonist-induced endocytosis of FPR2 as examined in several assay systems. Intriguingly, these data strongly indicate that FPR2 can undergo endocytosis through a β-arrestin/AP2-independent process. This is an internalization pattern shared with receptors for transferrin and endothelin-A, which are both endocytosed independently of β-arrestin and AP2, respectively [19]. Although Barbadin did not affect FPR2 internalization, it convincingly potentiated FPR2-mediated ROS production and promoted resensitization of desensitized FPR2s. A similar augmentation of the ROS response was also obtained at concentrations of FPR2 agonists (WKYMVM) that do not recruit β-arrestin or by FPR2 agonists (F2Pal_10_ and Cmp 14) that are very poor in recruiting β-arrestin, suggesting that this novel priming effect of Barbadin is achieved without β-arrestin recruitment. As mentioned above, the effect of Barbadin on FPR2-mediated ROS production resembled the effect induced by actin cytoskeleton-disrupting agents [11, 17, 18]. Despite this deviation from the prototypical mode of action for Barbadin, several lines of evidence suggest that Barbadin does not directly disrupt the actin cytoskeleton. These findings include that in contrast to latrunculin A, Barbadin had no effect on (i) the increase in F-actin polymerization induced by the FPR2 agonist WKYMVM, (ii) the actin cytoskeleton-dependent ROS production induced during phagocytic uptake of yeast particles was not affected by Barbadin, and (iii) the signals downstream of ATP-activated P2Y_2_Rs generated only when the actin cytoskeleton has been disrupted. Altogether, these observations indicate that Barbadin primes neutrophils and resensitizes/reactivates desensitized receptors through a mechanism resembling that of actin cytoskeleton-disruptive agents. However, as compared to actin cytoskeleton-disruptive agents, the effects exerted by Barbadin do not appear to involve a direct effect on the integrity of the actin cytoskeleton. At present, the precise mechanism underlying the influence of Barbadin on FPR2 activity is not known, but as Barbadin lacks effect on the FPR2-induced transient rise in [Ca^2+^]_i_, a general modulation of downstream signaling of agonist-occupied FPR2 is unlikely. Regarding assembly and activation of the electron-transporting NADPH-oxidase, a large number of stimuli (including many GPCR agonists) can induce ROS production in neutrophils, but not all signaling pathways that regulate these activation processes have been identified yet. However, it has been established that there is no direct link between GPCR-mediated activation of the oxidase and the transient rise in [Ca^2+^]_i_ [5, 18, 32, 39, 40], which gains further support from the data presented in this study. Hence, even although it is difficult to directly correlate NADPH-oxidase activity and calcium signaling during receptor reactivation induced by Barbadin or latrunculin A, our data corroborate previous studies demonstrating that a rise in [Ca^2+^]_i_ is not a requirement for activation of the NADPH-oxidase. Furthermore, our data support the notion that the neutrophil NADPH-oxidase can be activated in the absence of β- arrestin recruitment [9-11]. It follows that Barbadin is a biased and functionally selective regulator of FPR2 signaling as it influences ROS production by activating NADPH-oxidase without affecting calcium mobilization and neutrophil granule secretion.

Although β-arrestin modulated functions are inhibited by Barbadin, the AP2 inhibitor lacks direct effects on receptor-mediated β-arrestin recruitment [19]. Our earlier reports have demonstrated that FPR2 agonists that are potent stimuli in triggering calcium signaling, ERK1/2 phosphorylation and ROS production, but differ in their ability to recruit β-arrestin, also vary in their ability to induce neutrophils chemotaxis [9-11]. The present study shows that the functionally selective deviation linked to the ability to recruit β-arrestin is retained in the presence of Barbadin, further supporting the proposed mode of action of Barbadin in that it lacks a direct effect on receptor mediated β-arrestin recruitment [19]. The observation that Barbadin potentiates FPR2-mediated ROS production, no matter whether this was caused by the β-arrestin recruiting WKYMVM peptide or FPR2 agonists that are very poor in recruiting β-arrestin, suggests that the effects of Barbadin on FPR2-mediated ROS production is not dependent on β-arrestin. Several *in vitro* as well as *in vivo* processes potentiate FPR-mediated ROS production, and increased surface receptor expression as a result of granule secretion has been suggested as an important mechanism underlying the potentiation [17, 29, 30]. However, this is not the mechanism involved in the priming effect of Barbadin, demonstrated by its inability to induce the mobilization of granules. This conclusion is supported by the fact that while granule mobilization is an irreversible process the effects of Barbadin are reversible. Thus, the mechanism by which Barbadin potentiates FPR2-mediated ROS remains to be elucidated. Given that Barbadin binds to AP2 to prevent β-arrestin binding, the role of AP2 in the observed effects on ROS priming needs to be further investigated. With respect to the role of AP2, it is interesting to note that a comparison between EC_50_ values reveals that the potency of Barbadin mediated augmentation of ROS production in neutrophils is the same as that found for its inhibition of the β-arrestin-AP2 interaction. This suggests that the Barbadin-mediated effect on ROS production could be a result of its action on AP2 (this study and [19]). However, other target proteins including other binding partners for AP2, such as AP180, ARH and Scr [19] can, at this point, not be excluded until the modulating effect (if any) of Barbadin on these AP2 binding molecules has been be determined.

In summary, this study demonstrates some novel effects of the AP2 binding compound Barbadin. Although Barbadin did not affect agonist-induced endocytosis of FPR2, a process shown to be independent of whether the agonist recruits β-arrestin or not, Barbadin both increased FPR2 agonist induced ROS production, and resensitized agonist-desensitized FPR2 to produce ROS. Notably, these effects of Barbadin on FPR2 also proved independent of whether the agonist recruited β-arrestin or not. The effect of Barbadin on FPR2 induced neutrophil ROS production is very similar to the actin cytoskeleton-disrupting agent latrunculin A, albeit without altering other neutrophil functions regulated by a dynamic polymerization of the actin cytoskeleton. Elucidation of the precise mechanism(s) of Barbadin regarding its priming effect on the FPR2-mediated ROS production in neutrophils would lead to an increased understanding of the underlying molecular mechanisms regulating inflammatory reactions that are dependent on redox reactions. Barbadin and structurally related analogs of this AP2 inhibitor are expected to serve as useful molecular tools for further mechanistic studies of GPCR regulation in neutrophils.

## Abbreviations

Aoc: 2-aminooctanoic acid;
AF: Alexa Fluor;
AP2: clathrin adaptor protein 2;
AUC: area under the curve;
BRET: bioluminescence resonance energy transfer;
BSA: bovine serum albumin;
CL: chemiluminescence;
DMEM: Dulbecco’s modified Eagle medium;
DMSO: dimethyl sulfoxide;
DPBS: Dulbecco’s phosphate-buffered saline;
FPR: formyl peptide receptor;
GPCR: G protein-coupled receptor;
HBSS: Hank’s balanced salt solution;
HRP: horseradish peroxidase;
Lau: Lauroyl;
KRG: Krebs-Ringer phosphate buffer:
MOI: multiplicity of infection;
MPO: myeloperoxidase;
βNPhe: N-phenylmethyl- β-alanine;
βNrpe: N-(*R*)-1-phenylethyl-β-alanine;
PFA: paraformaldehyde;
PKC: protein kinase C;
PLC: phospholipase C;
PMA: phorbol 12-myristate 13-acetate;
RhB: rhodamine B;
RT: room temperature;
ROS: reactive oxygen species;
SOD: superoxide dismutase;
TR- FRET: time-resolved förster resonance energy transfer

## Acknowledgement

This work was supported by the Swedish Research Council (2015-005601; 2015-02448), the King Gustaf V 80-Year Foundation (FAI-2014-0011; FAI-2014-0029), Swedish foundation for strategic research (SM17-0046), the Magnus Bergwall foundation (MS2018-02579), the Swedish government under the ALF-agreement, Åke Wibergs Foundation (M15-0051), the Rådman and Mrs Ernst Collianders Foundation (MS 2019), the Ingabritt and Arne Lundberg Foundation and the European Union’s Horizon 2020 research and innovation programme (Marie Sklodowska-Curie grant agreement No 797497). S.C.W. is supported by a fellowship from the Swedish Society for Medical Research (P18-0098). M.B. is funded by the Canadian Institutes of Health Research (CIHR; FDN-148431) and holds a Canada Research Chair in Signal Transduction and Molecular Pharmacology. We thank Christian Le Gouill for plasmid DNA used in this study

## Authors’ contributions

H Forsman, CD and MS designed and oversaw all the aspects of the study. MS, AH, SCW, ES and TCM performed and analyzed the experiments with input from CD, KJ, MB and H Franzyk. H Forsman, MS, MB, and CD wrote the manuscript with all authors revising and approving it before submission of the final version.

